# CO_2_ Reduction by the Iron Nitrogenase Competes with N_2_ Fixation Under Physiological Conditions

**DOI:** 10.1101/2023.11.30.569367

**Authors:** Niels N. Oehlmann, Frederik V. Schmidt, Marcello Herzog, Annelise L. Goldman, Johannes G. Rebelein

**Affiliations:** Research Group Microbial Metalloenzymes, Max Planck Institute for Terrestrial Microbiology; Karl-von-Frisch-Straße 10, 35043 Marburg, Germany; Department of Plant and Microbial Biology, University of California Berkeley, Berkeley, California, 94720, USA

**Author notes:** Authors contributed equally (Niels N. Oehlmann, Frederik V. Schmidt).

## Abstract

Nitrogenases are the only known enzymes that reduce molecular nitrogen (N_2_) to ammonia. Recent findings have demonstrated that nitrogenases also reduce the greenhouse gas carbon dioxide (CO_2_), suggesting CO_2_ to be a competitor of N_2_. Intriguingly, nitrogenase isoforms (*i*.*e*., molybdenum (Mo), vanadium and iron (Fe) nitrogenase) differ significantly in their ability to reduce CO_2,_ but the mechanisms underlying these differences remain elusive. Here, we study the competing reduction of CO_2_ and N_2_ by the two nitrogenases of *Rhodobacter capsulatus*, the Mo and Fe nitrogenase. Analyzing their full CO_2_ reduction product spectrum *in vitro*, we find the Fe nitrogenase almost three-fold more efficient in CO_2_ reduction than the Mo isoform. Furthermore, the *in vitro* competition experiments reveal the Fe nitrogenase to be profoundly less selective for the reduction of N_2_ than the Mo nitrogenase. We observe the same effects *in vivo*, where adding CO_2_ drastically increases the doubling times of diazotrophically grown *R. capsulatus* strains that rely on the Fe nitrogenase. The Fe nitrogenase-dependent *R. capsulatus* strains reduce CO_2_ to methane under physiological conditions, highlighting the potential of the Fe nitrogenase for the biotechnological conversion of CO_2_ into value-added compounds. Furthermore, both products are secreted into the surrounding, potentially influencing the composition of microbial communities in Mo-deficient environments.

## 1. Introduction

Bioavailable nitrogen (N) is required to build life’s central metabolites, *e*.*g*. nucleotides and amino acids.^1, 2^ Although large parts of the Earth’s atmosphere consist of elemental dinitrogen (N_2_), the kinetic stability of N_2_ prevents its activation by most organisms. Only microorganisms expressing the enzyme nitrogenase can reduce N_2_ to ammonia (NH_3_); these microorganisms are called diazotrophs.^3^

Three nitrogenase isoforms are known in nature, which are distinguished by the heterometal content of their active site cofactor. The molybdenum (Mo) nitrogenase (encoded by *nifHDK*) is present in all diazotrophs and, thus, the most abundant isoform. In contrast, only a few diazotrophic microorganisms harbor vanadium (V, encoded by *vnfHDGK*) or iron (Fe, encoded by *anfHDGK*) nitrogenases.^4^ Mo nitrogenases have a higher efficiency for reducing N_2_ and are expressed in the presence of Mo. The alternative V or Fe nitrogenases are believed to function as fail-safe enzymes under Mo starvation.^5^ All three nitrogenases consist of a reductase component (NifH_2_, VnfH_2_, AnfH_2_) and a catalytic component (Nif(DK)_2_, Vnf(DGK)_2_, Anf(DGK)_2,_ Figure 1).^6-11^ The catalytic component consists of two symmetrical halves, each harboring a [Fe_8_S_7_]-cluster (P-cluster) and an active site cofactor. The latter is a [MFe_7_S_9_C-(R)-homocitrate]-cluster for the Mo and Fe nitrogenase (where M is either Mo or Fe) and a [VFe_7_S_8_C(CO_3_)-(R)-homocitrate]-cluster for the V nitrogenase. Based on the contained heterometal, the clusters are termed FeMoco, FeVco or FeFeco. The reductase component contains two adenosine triphosphate (ATP) binding sites and a [Fe_4_S_4_]-cluster at the protein’s homodimeric interface.

**Figure 1.**
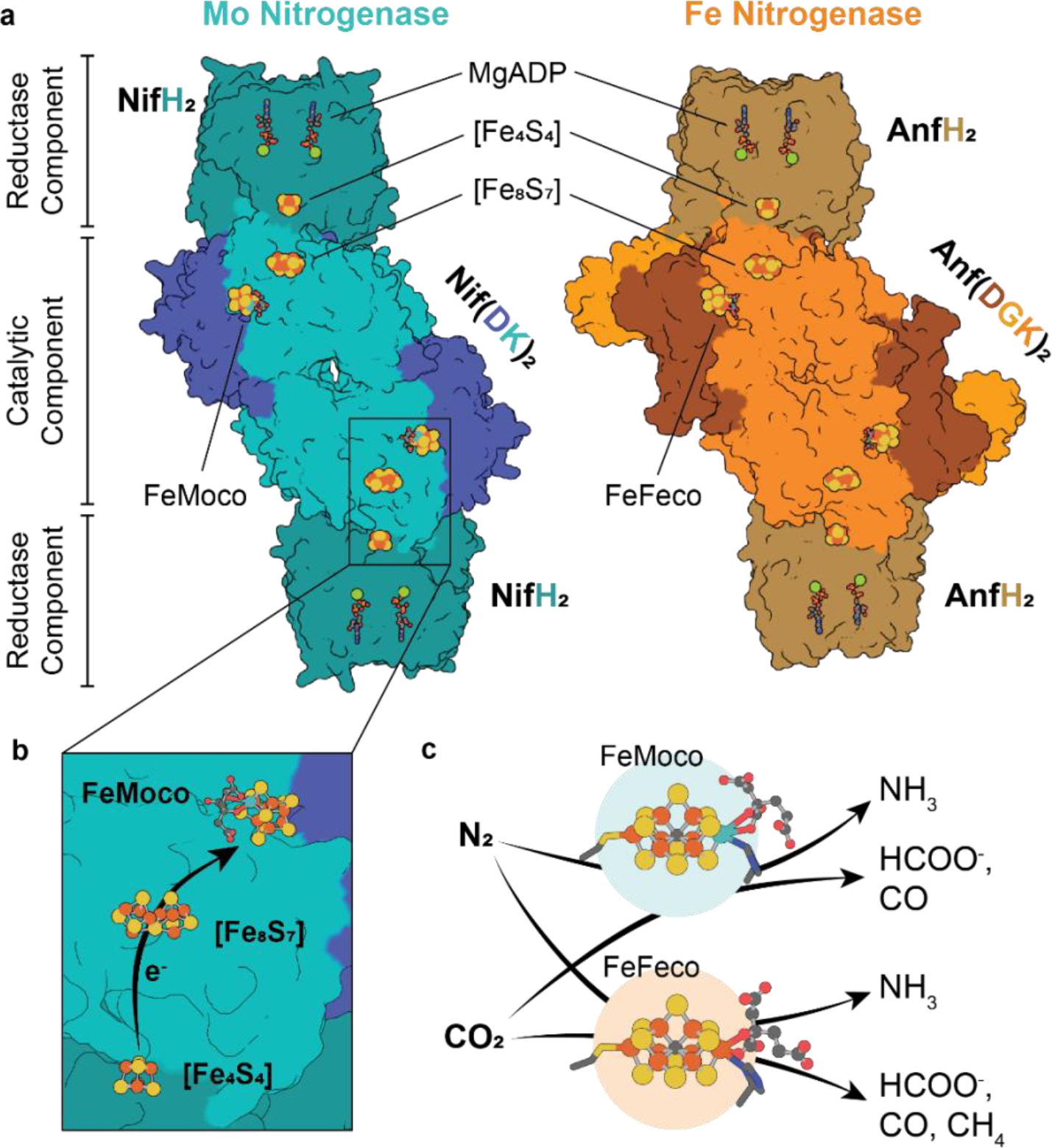
The nitrogenases of *Rhodobacter capsulatus* and their reactivities. (a) Architecture of the Mo (left, PDB: 7UTA) and Fe (right, PDB: 8OIE) nitrogenase. (b) Electron flow through the Mo nitrogenase. Electrons are delivered by the [Fe_4_S_4_] cluster of the reductase component and transferred via the [Fe_8_S_7_] cluster (P-cluster) to the FeMoco. The same scheme applies to the Fe nitrogenase. (c) Product spectrum of Mo (top) and Fe (bottom) nitrogenase for the reduction of N_2_ and CO_2_.

In the ATP-bound state, the reductase component transiently associates with the catalytic component, enabling the transfer of a low-potential electron from the [Fe_4_S_4_]-cluster via the P-cluster to the active site cofactor. Following the electron transfer, both molecules of ATP are hydrolyzed, and the reductase component dissociates from the catalytic component, allowing the cycle to repeat itself.^12, 13^ This way, the active site cofactor accumulates electrons, eventually used to reduce substrates.^3^ Despite the similar architecture, different reaction stoichiometries are observed for the reduction of N_2_ (at 1 atm) by the three nitrogenase isoforms (equations 1 – 3).^14^

Mo nitrogenase:

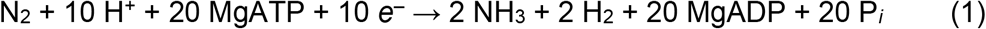

V nitrogenase:

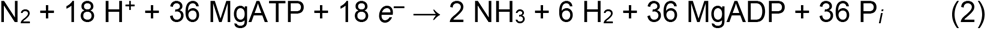

Fe nitrogenase:

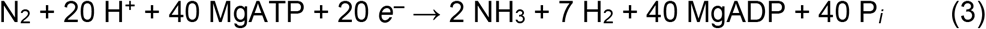

The N_2_ reduction requires a low reduction potential of the active site cofactors (*E*_*1/2*_ / V *vs*. normal hydrogen electrode FeVco –0.38 V; FeFeco –0.40 V).^15^ Thus, nitrogenases possess the power for reducing a variety of small molecules if they can reach the active site. Consequently, nitrogenases belong to the enzymes with the vastest known promiscuous activity.^4, 16^ For example, nitrogenases were shown to reduce non-physiological nitrogen compounds (azide, nitrite),^17-19^ carbon–nitrogen compounds (cyanide,^19^ (iso)nitriles),^20-22^ unsaturated hydrocarbon compounds (acetylene, cyclopropene),^23-27^ carbon monoxide^28-31^ and the potent greenhouse gas carbon dioxide (CO_2_).^32-35^

The reduction of CO_2_ by nitrogenase was first described for the Mo isoform, which converts CO_2_ to carbon monoxide (CO) at a rate of ∼0.09 nmol×(nmol Nif(DK)_2_×min)^−1^.^32^ Later, formate (HCOO^−^) was identified as the main product of the reaction, which is formed at 100-fold higher rates (9.8 nmol×(nmol Nif(DK)_2_×min)^−1^) than CO.^34^ More recently, also the alternative nitrogenases were found to reduce CO_2_. Intriguingly, the V nitrogenase performs carbon-carbon coupling reactions during CO_2_ reduction, releasing short-chain hydrocarbons like ethene and ethane besides CO.^30, 33^ The Fe nitrogenase was shown to convert CO_2_ to HCOO^−^ and methane (CH_4_).^35, 36^ Notably, the electron flux of the Fe nitrogenase towards the reduction of CO_2_ was found to be equivalent to the electron flux going into N_2_ reduction *in vitro*.^37^

CO_2_ is the primary waste product of modern fossil fuel-based economies and the greatest contributor to human-made climate change.^38^ Therefore, its removal and utilization from the atmosphere is a major societal challenge driving the need for CO_2_ reduction processes to establish a carbon-neutral economy.^39^ Given their unique reactivities, nitrogenases might offer new approaches to convert CO_2_ directly into biofuels and feedstock chemicals like methane, ethene or propene.^31^ However, the impact of CO_2_ on nitrogenase activity and the growth of diazotrophs remains elusive. Since CO_2_ is a common metabolite present in all living cells,^40^ nitrogenases are constantly exposed to CO_2_ that might compete with the reduction of N_2_ (Figure 1c).

Here, we study the competing reduction of N_2_ and CO_2_ by the Mo and the Fe nitrogenase of *Rhodobacter capsulatus*. Following the purification of both nitrogenases, we characterized their full CO_2_ product spectrum *in vitro* and compared their electron flux under argon (Ar), N_2_ and CO_2_ atmospheres. We find significant differences between the two isozymes, with the Mo nitrogenase being very selective for N_2_ fixation and the Fe nitrogenase being more promiscuous towards CO_2_ reduction. We observe the same effect *in vivo*, where the promiscuity of the Fe nitrogenase leads to a CO_2_-dependent deceleration of the diazotrophic growth of *R. capsulatus* strains relying on the Fe nitrogenase. The Fe nitrogenase expressing strain reduces intracellularly produced CO_2_ to CH_4_ and HCOO^−^ suggesting that the Fe nitrogenase functions as a N_2_ and CO_2_ reductase under physiological conditions.

## 2. Results and Discussion

To comprehensively characterize the reduction of CO_2_ by the Mo and Fe nitrogenase we used our recently established plasmid-based expression and purification of the Fe nitrogenase in *R. capsulatus*.^11^ We extended the system by establishing the recombinant production and purification of the Mo nitrogenase (see Materials & Methods section). After purifying both nitrogenase enzymes, we confirmed their full H^+^ and N_2_ reduction activity *in vitro* (Figure S1). The nitrogenase-specific activity followed the expected hyperbolic trend for dihydrogen (H_2_) and NH_3_ formation under an increasing ratio of reductase to catalytic component.^41^ Both isozymes reached the activity plateau at the expected reductase component excess (Mo nitrogenase: ∼20:1, Fe nitrogenase: ∼40:1). They exhibited the previously observed NH_3_ formation rates under high electron flux conditions (Mo nitrogenase: 107 ± 6 nmol×(nmol Nif(DK)_2_×min)^−1^, Fe nitrogenase 36.4 ± 1.0 nmol×(nmol Anf(DGK)_2_×min)^−1^, Figure S1).^42^

After confirming that the purified Mo and Fe nitrogenase were fully active for N_2_ and H^+^ reduction, we proceeded by analyzing their ability to reduce CO_2_ *in vitro*. (Figure 2). Under a CO_2_ atmosphere, the Mo nitrogenase formed mainly H_2_, followed by HCOO^−^ with formation rates of 168.7 ± 1.6 nmol×(nmol Nif(DK)_2_×min)^−1^ and 58 ± 2 nmol×(nmol Nif(DK)_2_×min)^−1^, respectively (Figure 2a). Besides H_2_ and HCOO^−^, the Mo nitrogenase also formed traces of CO at a rate of 0.40 ± 0.03 nmol×(nmol Nif(DK)_2_×min)^−1^, but we could not detect any hydrocarbons (Figure 2b). In contrast, the main product of the Fe nitrogenase was found to be HCOO^−^ (60 ± 5 nmol×(nmol Anf(DGK)_2_×min)^−1^), followed by H_2_ (34 ± 6 nmol×(nmol Anf(DGK)_2_×min)^−1^, Figure 2c). In addition, we found the Fe nitrogenase produced CO and CH_4_ at rates of 0.19 ± 0.03 nmol×(nmol Anf(DGK)_2_×min)^−1^ and 0.026 ± 0.003 nmol×(nmol Anf(DGK)_2_×min)^−1^, respectively (Figure 2d). The higher obtained HCOO^−^ formation activity (58 ± 2 nmol×(nmol Nif(DK)_2_×min)^−1^) for the Mo nitrogenase of *R. capsulatus* and the reported value (9.8 nmol×(nmol Nif(DK)_2_×min)^−1^) for the Mo nitrogenase of *A. vinelandii* is likely due to the higher partial pressure of CO_2_ (1.2 atm *vs*. 0.45 atm) used in our experiments.^34^ Interestingly, both, the Fe and Mo nitrogenase exhibit the same specific activity for HCOO^−^ formation (∼60 nmol×(nmol-×min)^−1^) despite the more than two-fold higher electron flux of the Mo nitrogenase. The same HCOO^−^-formation rate could indicate a common rate-limiting step for both nitrogenases in reducing CO_2_ to HCOO^−^, independent of the electron flux. Moreover, the ratio of reductase to catalytic component of the Fe nitrogenase required for the high electron flux reduction decreased from ∼40:1 for N_2_ reduction to ∼20:1 for CO_2_ reduction (Figure 2c, d). This result might indicate distinct rate-limiting steps for CO_2_ and N_2_ reduction of the Mo and Fe nitrogenase. For N_2_ reduction under high electron flux the hydrolysis of ATP and thus the release of the reductase component is regarded to be the rate limiting step.^43^ Since we observe lower reductase to catalytic component ratios to be sufficient for maximal activity for HCOO^−^ formation by nitrogenases, a step different from ATP hydrolysis might be rate limiting for CO_2_ reduction.

**Figure 2.**
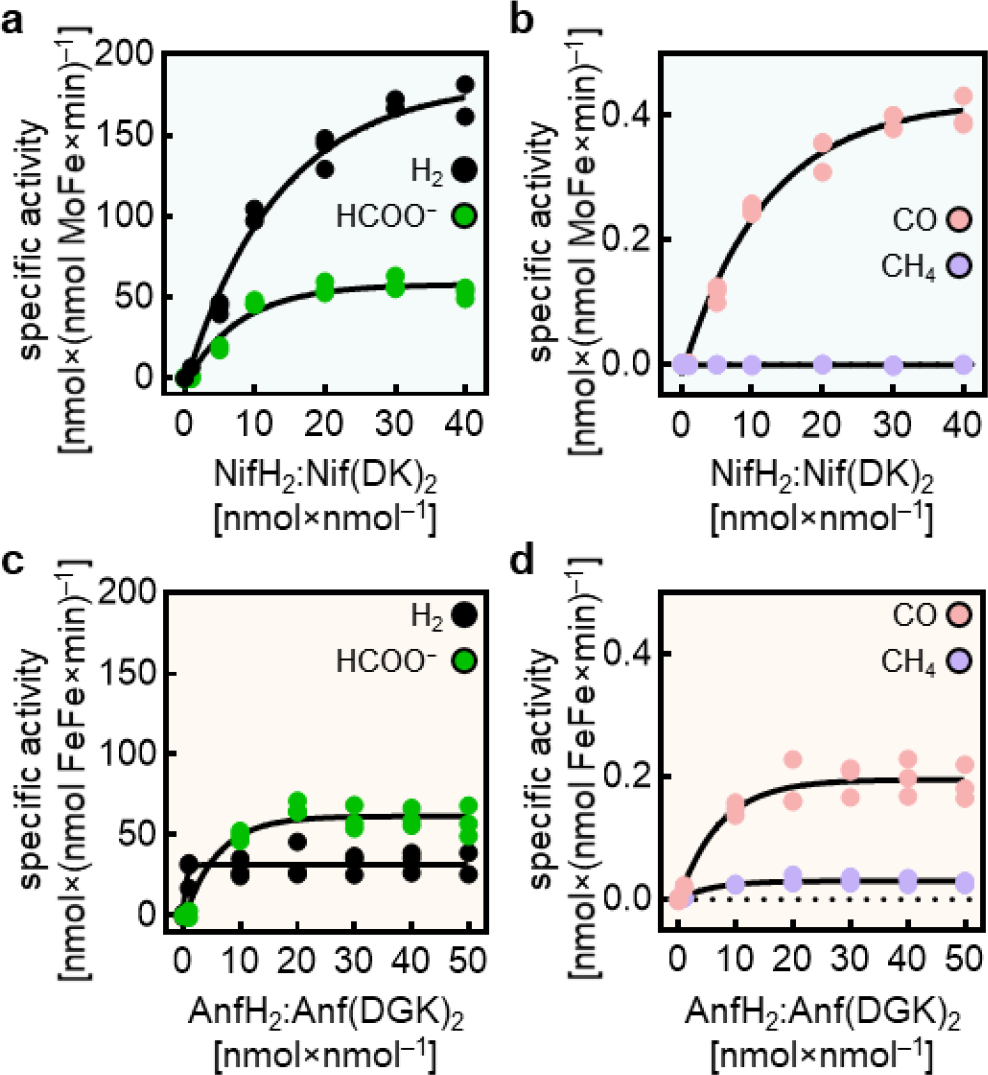
*In vitro* CO_2_ reduction by the Mo and Fe nitrogenase. Product formation rates of the Mo (a, b) and Fe nitrogenase (c, d) for *in vitro* activity assays conducted under a 1.2 atm CO_2_ atmosphere. Plotted are specific activities for the formation of H_2_ and HCOO^−^ (a, c) and CO and CH_4_ (b, d) versus the molar excess of reductase component over catalytic component (n = 3).

In conclusion, the Mo and Fe nitrogenase show distinct product profiles for the reduction of CO_2_. In contrast to the Mo isoform, the Fe nitrogenase can reduce CO_2_ directly to CH_4_ and uses most electrons to form HCOO^−^ while the Mo nitrogenase mainly forms H_2._ The rate of HCOO^−^ formation is similar for both nitrogenase enzymes and seems, surprisingly, to be independent of the electron flux.

To gain a deeper understanding of the observed activities, we continued by analyzing the overall electron flux under Ar, N_2_ and CO_2_ *in vitro* to evaluate the efficiency of the Mo and Fe nitrogenase in directing electrons into the different products, following the approach of the nitrogenase field (Figure 3a, b, e, f).^34, 36^ Under an Ar atmosphere, the Mo nitrogenase shows a more than two-fold higher electron flux towards the formation of H_2_ than the Fe isoform. Changing the headspace to N_2_, we observed a drop in the Mo nitrogenase electron flux by 47%, whereas the overall electron flux of the Fe nitrogenase dropped by only 17%. This result follows previously observed trends for the nitrogenase enzymes from *Azotobacter vinelandii*.^44^ Similarly, when changing the reaction atmosphere from Ar to CO_2_, we also observed a drop in the overall electron flux for both isoenzymes (Mo nitrogenase: 60% drop; Fe nitrogenase: 42% drop; Figure 3a, e). Intriguingly, previous studies have demonstrated that the Mo nitrogenase electron flux does not change between Ar and N_2_ atmospheres in a cyclic voltammetry setup that includes electron mediators like methyl viologen.^45^ This implies that unaccounted reduction reactions cause the apparent drop in electron flux by switching from Ar to N_2_. Recent work by Tanifuji *et al*. suggested that releasing NH_3_ from the active site cofactor of the Mo nitrogenase requires replenishment of the belt sulfides by reducing 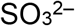 thereby causing a previously unaccounted electron flux.^46^ Thus, the observed drop of electron flux for the two isozymes when switching to a CO_2_ atmosphere indicates that this mechanism could also apply to the CO_2_ reduction reaction and might imply that CO_2_ has a similar binding mode as N_2_.

**Figure 3.**
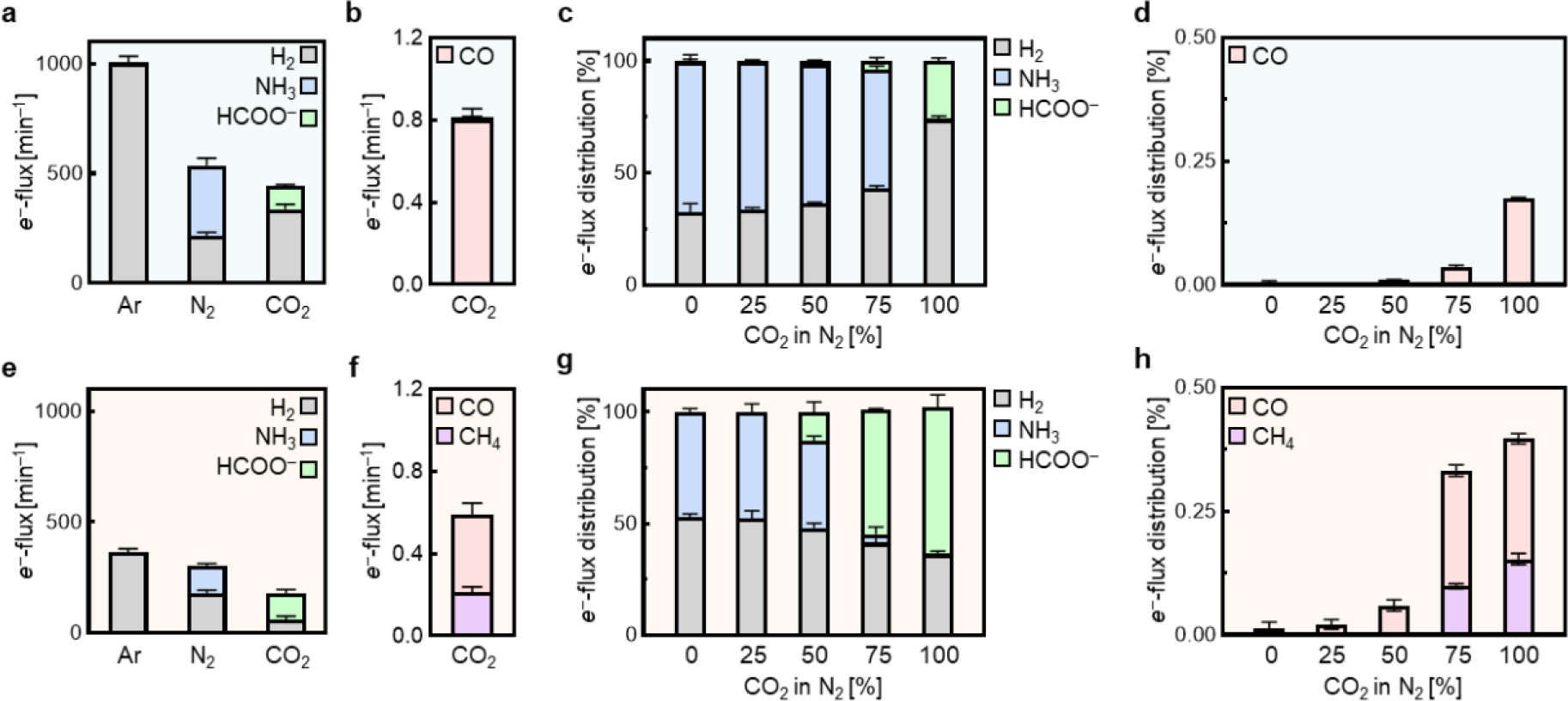
Electron flux analysis of the Mo and Fe nitrogenase. Left: Electron flux towards the formation of H_2_, NH_3_, HCOO^−^, CO and CH_4_ *in vitro* under Ar, N_2_ and CO_2_ atmospheres for the Mo nitrogenase (**a, b**) and Fe nitrogenase (**e, f**). Right: Electron flux distribution of *in vitro* competition assays of Mo (**c, d**) and Fe nitrogenase (**g, h**) under mixed atmospheres of CO_2_ and N_2_. Bars and error bars represent mean ± standard deviation from replicates (n = 3).

We also found that the flux of electrons towards H_2_ formation increased for the Mo nitrogenase while being suppressed for the Fe nitrogenase when we changed the headspace gas from N_2_ to CO_2_ (Figure 3a, e). This resulted in the Fe nitrogenase having a 2.75-fold higher efficiency for CO_2_ reduction, with 66% of the total electron flow ending in CO_2_ reduction products in the case of the Fe nitrogenase and only 24% for the Mo nitrogenase. Remarkably, the Fe nitrogenase exhibited a similar electron flux towards HCOO^−^ formation (116 ± 9 *e*^−^×min^−1^) under a CO_2_ atmosphere as towards NH_3_ (124 ± 5 *e*^−^×min^−1^) under a N_2_ atmosphere, which emphasizes the ability of the Fe nitrogenase to function as a N_2_ and a CO_2_ reductase (Figure 3e). In conclusion, the Fe nitrogenase appears to be more efficient in CO_2_ reduction than the Mo nitrogenase.

Intrigued by the similar activity of the Fe nitrogenase for CO_2_ and N_2_ reduction, we wondered whether CO_2_ is a competitor for N_2_ reduction for the Fe nitrogenase and investigated the substrate specificity in more detail. For this purpose, we prepared *in vitro* activity assays under mixed atmospheres of CO_2_ and N_2_ and compared the Mo and Fe nitrogenase electron flux for the individual reduction products (Figure 3c, d, g, h). For the Mo nitrogenase, the electron flux towards NH_3_ formation decreased only by 10% when the CO_2_ concentration was increased from 0% to 75% (Figure 3c). Consequently, NH_3_ remains the main product in all assays where N_2_ is present in the headspace. HCOO^−^ formation was first observed under an atmosphere of 75% CO_2_ in N_2_ (3.7% of total electron flux) and increased to 25.8% of the total electron flux under 100% CO_2_. We observed the same trend for the electron flux towards CO formation that started at 75% CO_2_ and increased five-fold in the absence of N_2_ (Figure 3d). In contrast to the Mo isoform, the Fe nitrogenase electron flux towards NH_3_ formation decreased dramatically from 50% in the absence of CO_2_ to only 6% under 75% CO_2_ (Figure 3g). Moreover, the formation of HCOO^−^ by the Fe nitrogenase already started at 50% CO_2_ and HCOO^−^ became the main product of the reaction under an atmosphere of 75% CO_2_. Formation rates of CO and CH_4_ increased accordingly with rising concentrations of CO_2_ in the headspace (Figure 3h). Notably, we observed similar trends for both nitrogenases in CO_2_ titration experiments conducted under an Ar atmosphere. However, the onset of CO_2_ reduction was earlier (50% CO_2_ for the Mo nitrogenase and 25 % CO_2_ for the Fe nitrogenase) compared to the results obtained in N_2_ (Figure S2). In conclusion, the Mo nitrogenase appears to be more selective for reducing N_2_ and the Fe nitrogenase is more promiscuous for reducing CO_2_ even in the presence of N_2_.

Despite the difference in selectivity, the Mo nitrogenase readily reduces CO_2_ in the absence of N_2_, leading us to speculate that a structural feature of the Mo nitrogenase (*i*.*e*., the substrate channel) is responsible for the discrimination of CO_2_ in the presence of N_2_. To investigate this hypothesis, we compared the substrate channels of the Mo and Fe nitrogenase. The likeliest substrate channel of the Mo nitrogenase is the Igarashi-Seefeldt (IS) channel,^47^ which has been characterized through molecular dynamics simulations in combination with Fourier-transformed infrared spectroscopy.^48^ The IS channel comprises 15 residues, all located within the NifD subunit, forming a hydrophobic channel from the nitrogenase surface to the FeMoco (Figure S3a-d). Aligning the Mo nitrogenase structure with the recently solved Fe nitrogenase structure, we could identify a homologous channel in AnfD (Figure S3e-h), which CAVER predicted to be the best substrate channel of the Fe nitrogenase.^11^ Intriguingly, we found only four of the 15 channel residues to be conserved in the Fe nitrogenase. While the altered amino acid residues do not change the hydrophobic character of the channel, they do reshape the channel and could influence the substrate’s access to the active site. Thus, the distinct substrate channels of the Mo and Fe nitrogenase might explain the observed differences in N_2_ *versus* CO_2_ selectivity.

Next, we tested whether the different selectivity of the two nitrogenases would affect the diazotrophic growth of *R. capsulatus* in the presence of CO_2_. For this purpose, we constructed *R. capsulatus* strains depending either on the Mo nitrogenase (MM0335) or the Fe nitrogenase (MM0057, Δ*nifD*::SpR, Δ*modABC*) and characterized their growth under an N_2_ atmosphere with increasing CO_2_ concentrations (Figure 4). In the absence of CO_2_, growth of *R. capsulatus* strains depending on the Mo nitrogenase showed a 23% lower doubling time (*T*_d_) than the Fe nitrogenase-dependent strain (Mo: *T*_d_= 6.65 ± 0.19 h *vs*. Fe: 8.6 ± 0.2 h), which is in line with the higher N_2_ fixing activity and efficiency of the Mo nitrogenase (compare Figure 4a, b to Figure 4c, d).^49,50^ Increasing the CO_2_ concentration in the culture headspace gradually to 20% CO_2_ caused the *T*_d_ of the Mo nitrogenase-dependent strain to increase slightly (12%) to 7.42 ± 0.1 h (Figure 4b). In contrast, the Fe nitrogenase-dependent strains exhibited a clear dose-response to rising concentrations of CO_2_. The *T*_d_ of the Fe nitrogenase expressing strain increased strongly already under 10% CO_2_ and reached a *T*_d_ of 15.0 ± 0.5 h (a 74% increase) under 20% CO_2_ (Figure 4d). In conclusion, our findings suggest that CO_2_ is a competitor to N_2_ for *in vivo* nitrogenase activity and appears to affect diazotrophic growth for Fe nitrogenase dependent strains.

**Figure 4.**
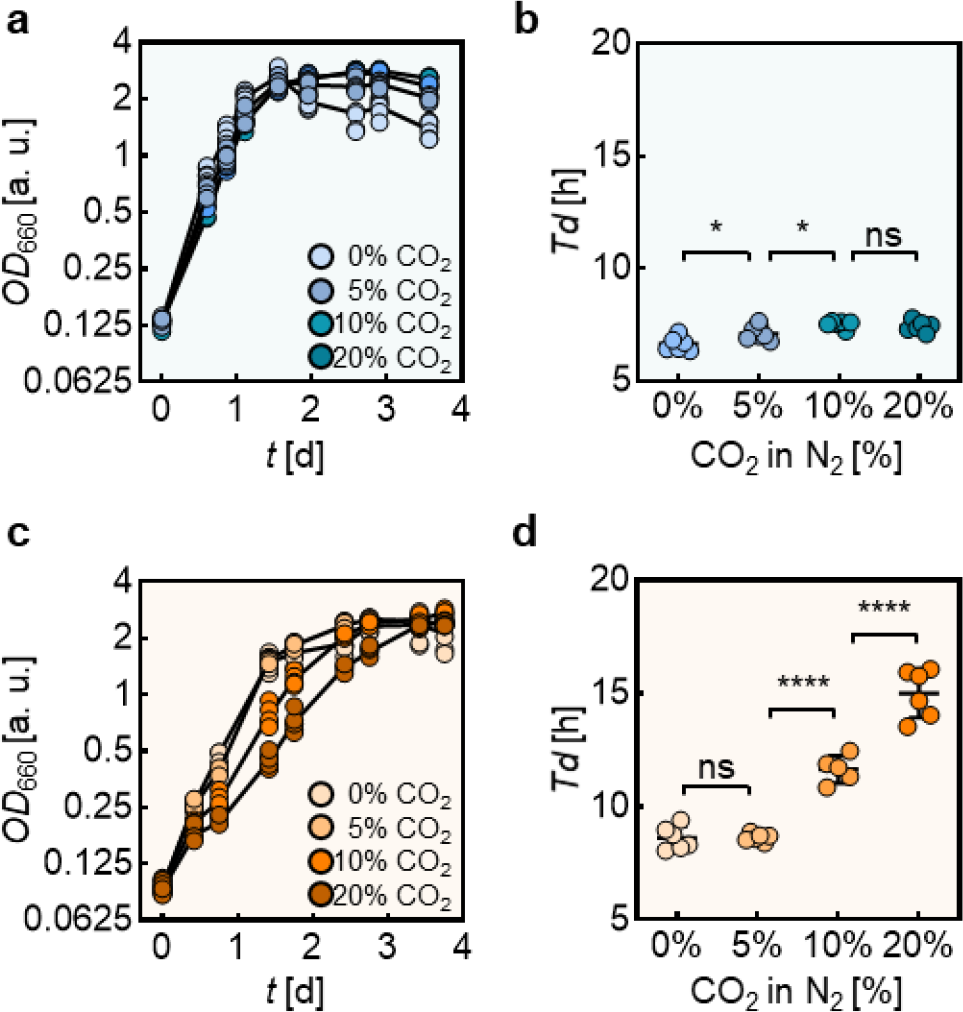
Effect of CO_2_ on diazotrophic growth of *R. capsulatus*. Diazotrophic growth curves *of R. capsulatus* strains expressing the Mo (**a**) and Fe (**c**) nitrogenases with increasing CO_2_ concentrations. (**b, d**) Corresponding doubling times (*T*_d_) in the exponential growth phase. Spheres represent the values from biological replicates (n = 6).

Based on our *in vitro* results, we hypothesized that the strong inhibitory effect of CO_2_ on the diazotrophic growth of Fe nitrogenase-dependent strains might arise from the high promiscuity of the Fe nitrogenase for the reduction of CO_2_. To address this hypothesis, we first monitored CO_2_ reduction products (HCOO^−^ and CH_4_) during diazotrophic growth in the absence and presence of 20% CO_2_ (Figure 5a, e). As demonstrated in our *in vitro* data, HCOO^−^ is the main product of CO_2_ reduction by nitrogenases. Due to its membrane permeability, it is released into the culture medium,^51^ allowing for easy quantification of metabolically derived HCOO^−^ and thus nitrogenase CO_2_-reducing activity. To account for non-nitrogenase-derived HCOO^−^ we compared the Mo and Fe nitrogenase-dependent strains to controls supplemented with NH_3_, which suppresses nitrogenase expression. Intriguingly, only the Fe nitrogenase-dependent strain exhibited significantly elevated HCOO^−^ concentrations in the culture medium compared to the NH_3_ controls (Figure 5a). Although Mo nitrogenase-derived HCOO^−^ formation is possible, the difference under Mo nitrogenase expressing and repressed conditions is insignificant. The addition of 20% CO_2_ to the culture headspace caused an increase in HCOO^−^ concentrations by 3-fold (4.54 ± 0.13 mM) for the Fe nitrogenase strains (Figure 5a). Since HCOO^−^ is secreted into the extracellular space, this reaction might impact the composition of bacterial communities as it provides an accessible C1 compound for other microorganisms. Notably, the HCOO^−^ concentrations increased during exponential growth before reaching a plateau after four days for the Fe nitrogenase strain (Figure S4). Thus, the onset of the plateau coincided with the cultures entering the stationary phase, which resembles the pattern known from N_2_ fixation by nitrogenases.^52, 53^ To exclude the interference of the formate dehydrogenase (FDH) with our results, we repeated the same experiments with FDH-deficient strains (Figure S5). The deletion of FDH did not change the previously obtained results. This led to our conclusion that the observed differences in HCOO^−^ formation were solely based on the respective nitrogenase isoform.

**Figure 5.**
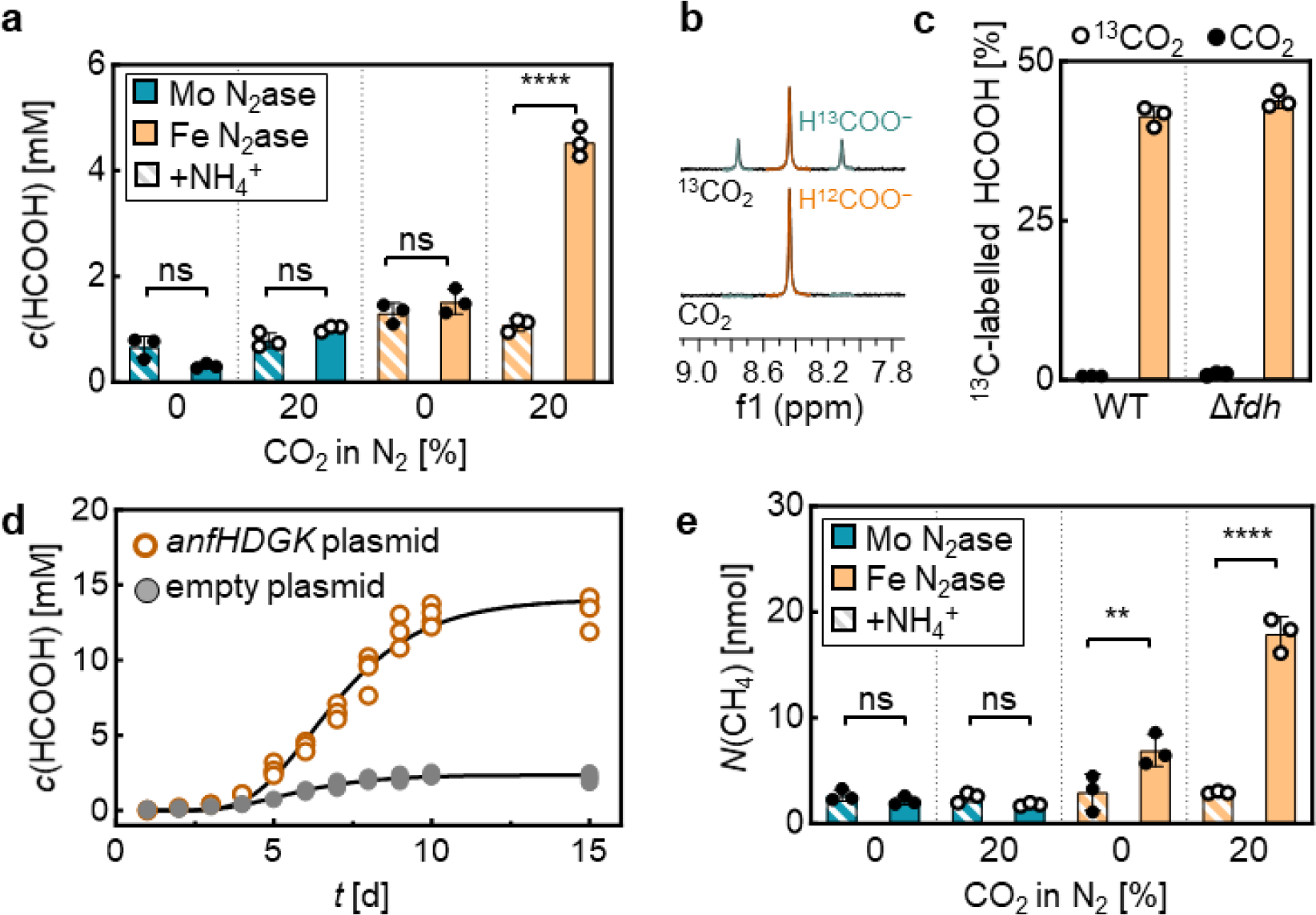
*In vivo* CO_2_ reduction by nitrogenase expressing *R. capsulatus* strains. (**a**) HCOO^−^ concentration in *R. capsulatus* culture supernatant expressing the Mo nitrogenase or the Fe nitrogenase under diazotrophic growth conditions in the absence or presence of 20% CO_2_ after 6 d of growth. (**b**) Representative ^1^H NMR (300.0 MHz, D_2_O) spectra of *R. capsulatus* culture supernatant incubated under 20% CO_2_ or ^13^CO_2_. Shown are the resonance peaks of H^12^COO^−^ (red) and H^13^COO^−^ (teal). (**c**) Fraction of ^13^C labelled HCOO^−^ in the culture supernatant of the Fe nitrogenase strain and the respective Δ*fdh strain* grown with 20% CO_2_ (black) or 20% ^13^CO_2_ (99%-^13^C enriched, white) in the culture headspace. The ratio between ^12^C- and ^13^C-labelled HCOO^−^ was determined by NMR spectroscopy. (**d**) HCOO^−^ accumulation over time in the culture medium of *R. capsulatus* Δ*anfHDGK* strain expressing the Fe nitrogenase from a plasmid (orange) or carrying an empty plasmid (grey). The cultures were cultivated under an atmosphere of 8% CO_2_ in Ar using glutamate as the N source. (**e**) Amount of CH_4_ measured in the culture headspace of *R. capsulatus* cells after 9 d of growth determined by GC-FID. Dots represent the individual values from biological replicates (**a-e**: n = 3).

To confirm that CO_2_ is the precursor for the observed reactions, we performed ^13^CO_2_ labeling experiments (Figure 5b, c). For this purpose, the Fe nitrogenase *R. capsulatus* strains with and without the FDH gene (WT or Δ*fdh*) were cultivated under an atmosphere of N_2_, supplemented with 20% CO_2_ or 20% ^13^CO_2_. After six days, extracellular HCOO^−^ was quantified by ^1^H NMR spectroscopy using the characteristic down field shifted signal at *δ* 8.44 ppm (Figure 5b). Here, the H^13^COO^−^ signal is split by ^1^*J*(^13^C,^1^H) coupling, which enables the relative quantification of the H^13^COO^−^ and H^12^COO^−^signal intensity. Both Fe nitrogenase-dependent strains showed high ^13^C-labelling ratios of 41.4% for the WT and 44.0% for the Δ*fdh* strain. No ^13^C enrichment between WT and *fdh* knockout is visible because the Fe nitrogenase-dependent strain features a genetic knockout of the molybdenum transporter (ModABC). This modification is necessary for producing the Fe nitrogenase because Mo is a potent transcriptional repressor of the Fe nitrogenase gene cluster. Since FDH is a Mo-dependent enzyme, the Fe nitrogenase strain grown without Fe shows no FDH-mediated CO_2_ and HCOO^−^ interconversion. Unlabeled HCOO^−^ likely originates from metabolically derived CO_2_ sufficient to fuel nitrogenase-mediated CO_2_ reduction. In summary, these results confirm CO_2_ as the origin of nitrogenase-derived HCOO^−^.

To unambiguously confirm that the Fe nitrogenase is responsible for the *in vivo* reduction of CO_2_, we created knockout strains, lacking both nitrogenase systems (*nifDK* and *anfHDGK*) as well as *fdh*. Next, we complemented the strain either with an *anfHDGK* expression plasmid or an empty control plasmid and supplemented the growth media with glutamate as the N source. Growing these two strains under an atmosphere of 8% CO_2_ in Ar, cultures of the *anfHDGK* complemented strain produced high amounts of HCOO^−^ (13.3 ± 0.4 mM). In contrast, the control strain with an empty plasmid produced only 2.18 ± 0.12 mM HCOO^−^ under identical conditions (Figure 5d). Thus, we conclude that the vast majority of extracellular HCOO^−^ originates from the Fe nitrogenase.

Analogously to the formation of HCOO^−^, we observed a CO_2_ concentration-dependent formation of CH_4_ that we quantified from the culture headspace (Figure 5e). Notably, we could only detect CH_4_ in the headspace of the Fe nitrogenase but not in the Mo nitrogenase strains, which stands in accordance with our *in vitro* results and previous literature.^35^ We could also detect significant amounts of CH_4_, beyond the background of the nitrogenase repressed culture, without the addition of any CO_2._ This observation indicates that the metabolically produced CO_2_ is directly converted by the Fe nitrogenase into CH_4_. Thus, we expect physiological CO_2_ concentrations to be sufficient for the formation of CH_4_ and HCOO^−^ in Fe nitrogenase expressing diazotrophs, also in the presence of N_2_.

## Conclusion

The recent findings on nitrogenases reducing CO_2_ suggest novel pathways for recycling carbon waste from the atmosphere.^32, 34, 54^ Nitrogenases’ ability to convert CO and CO_2_ into reduced hydrocarbons has been at the center of attention.^28-31, 33, 55-57^ At the same time, these findings challenge our overall understanding of nitrogenases as solely N_2_- and H^+^-reducing enzymes by suggesting CO_2_ to be a competing substrate for biological nitrogen fixation. This effect might be most relevant for alternative nitrogenases, which appear to have higher activities for the reduction of CO_2_ compared to the Mo isoform. Here, we analyzed the electron flux by the Mo and Fe nitrogenases under changing atmosphere compositions, *i*.*e*., Ar, N_2_ and CO_2_. We observe a drop in electron flux by both nitrogenases when changing the reaction atmosphere from Ar to N_2_ or CO_2_, which suggests mechanistic similarities between the reduction of N_2_ and CO_2_. Thus, we hypothesize that CO_2_ binds at the E2-E4 state of the Lowe-Thorneley model as proposed for N_2_ reduction.

Next, we analyzed the competing reduction of N_2_ and CO_2_ by the Mo and Fe nitrogenase from *R. capsulatus*. Under CO_2_, we find both enzymes to generate HCOO^−^ at similar rates, a remarkable result considering the two-fold higher overall electron flux of the Mo over the Fe nitrogenase.^44^ Our results demonstrate the Fe nitrogenase directs two-thirds of its electron flux into the reduction of CO_2_. This exceeds the electron flux observed for the formation of NH_3_ under an N_2_ atmosphere, leading to our conclusion that the Fe nitrogenase acts as an N_2_ and a CO_2_ reductase, simultaneously. In support of this hypothesis, the Fe nitrogenase reduces more CO_2_ than N_2_ in an atmosphere of 75% CO_2_ and 25% N_2_. In contrast, the Mo isoform is highly selective for N_2_ and only reduces CO_2_ at similar rates as the Fe nitrogenase in the absence of N_2_. Thus, we conclude that both substrates access the active site through the same substrate channel, which is more selective for N_2_ in the case of the Mo nitrogenase compared to the Fe nitrogenase. The lower selectivity of the Fe nitrogenase seems to be physiologically relevant, as we observe CO_2_-dependent deceleration of diazotrophic growth in *R. capsulatus* strains relying on the Fe nitrogenase. Interestingly, metabolically derived CO_2_ is sufficient to drive the Fe nitrogenase-catalyzed CH_4_ formation *in vivo* (Figure 5b), implying that this process is widespread in nature. Thus, the Fe nitrogenase-catalyzed reduction of CO_2_ might contribute to releasing the potent greenhouse gas CH_4_. Since HCOO^−^ and CH_4_ are secreted into the extracellular space, Fe nitrogenase activity provides C1 compounds to the surrounding microenvironment, thereby potentially influencing local microbiota to an unknown degree. In the absence of N_2_, we observed an accumulation of more than 13 mM HCOO^−^ in the culture supernatant of Fe nitrogenase expressing *R. capsulatus* strains. This ability stands out from other formate-producing enzymes like FDH or pyruvate formate lyase. FDH catalyzes a reversible reaction that is limited by the change in Gibbs energy for the formation of HCOO^−^ from CO_2_ and NADH to around 10 μM HCOO^−^ (pH 6.8, ionic strength of 0.25 M, c(CO_2_) = 2.86 mM, concentrations of other reactants set to 1 mM).^58^ In contrast, the Fe nitrogenase reduces CO_2_ irreversibly, achieved by coupling the reaction to ATP hydrolysis.

Our model organism, *R. capsulatus*, is a phototrophic organism that can produce ATP from light energy. Hence, *R. capsulatus* or similar phototrophic organisms could serve as a chassis for the sustainable, light-driven *in vivo* reduction of CO_2_ by the Fe nitrogenase, opening novel pathways for converting carbon waste into value-added compounds. Improving nitrogenases’ ability to reduce CO_2_ directly to CH_4_, C_2_H_4_ and C_2_H_6_ would allow mkithe carbon-neutral synthesis of fuels and bulk chemicals. Thus, nitrogenases potentially offer new solutions for transitioning into a circular economy.

## Acknowledgment

J.G.R. thanks the Deutsche Foschungsgemeinschaft (DFG, German Research Foundation) – 446841743 for funding. N.N.O. thanks the Fonds der Chemischen Industrie for a Kekulé fellowship. Furthermore, the authors thank the Rebelein laboratory for fruitful discussions and valuable comments on the manuscript. We thank B. Masepohl and T. Drepper for providing strains and plasmids.

## Author contributions statement

These authors contributed equally: Niels N. Oehlmann, Frederik V. Schmidt J.G.R. conceived and supervised the project. J.G.R. acquired funding. N. N. O., F.V.S. and J.G.R. designed and analysed experiments. F.V.S. and N.N.O. performed molecular work. F.V.S. and M.H. performed anaerobic protein purification. N.N.O., F.V.S. performed *in vitro* enzyme biochemistry. N.N.O. and A.L.G. performed *in vivo* experiments. N.N.O. performed CO_2_ labelling experiments. N. N. O., F.V.S. and J.G.R. wrote the original manuscript, which was reviewed and edited by all authors.

