## Supplemental Information for "CO_2_ Reduction by the Iron Nitrogenase Competes with N_2_ Fixation Under Physiological Conditions"

[†]Authors contributed equally.

**Safety Statement:** No unexpected or unusually high safety hazards were encountered.

### 1. General aspects

Unless noted otherwise, all chemicals were purchased from Carl Roth GmbH + Co. KG (Karlsruhe, Germany), Thermo Fisher Scientific Inc. (Waltham, USA), Sigma-Aldrich (St. Louis, USA) or Tokyo Chemical Industry Deutschland GmbH (Eschborn, Germany) and were used directly without further purification. Gases were purchased from Air Liquide Deutschland GmbH (Düsseldorf, Germany)

All primers used in this study were purchased from Eurofins Genomics (Ebersberg, Germany) and are listed in Table 3. All plasmids used and created in this study are listed in Table 2. Polymerase chain reactions (PCRs) were conducted with Q5® High-Fidelity DNA Polymerase (New England Biolabs, Ipswich, USA), PCR purifications with the Monarch® PCR & DNA cleanup kit (New England Biolabs, Ipswich, USA), extraction of genomic DNA with the Monarch® Genomic DNA Purification Kit (New England Biolabs, Ipswich, USA), Gibson assemblies with the NEBuilder® HiFi DNA Assembly Master Mix (New England Biolabs, Ipswich, USA), restriction digest ligation cloning with T4 DNA Ligase (New England Biolabs, Ipswich, USA) and Golden Gate cloning with the NEBridge® Golden Gate Assembly Kit (New England Biolabs, Ipswich, USA) according to the instructions provided by the manufacturer. Successful assembly of desired vectors was verified by Sanger sequencing through Microsynth Seqlab GmbH (Göttingen, Germany).

### 2. Molecular cloning

For the generation of plasmid pMM0156, pK18mobSacB was linearized by PCR using Primers P1 and P2. The *nifHDK* operon was PCR amplified from *R. capsulatus* genomic DNA (Strain B10S) with P3 and P4. The purified fragments were cloned via Gibson assembly and the reaction mix was used to transform chemo competent DH5α cells. The outgrowth was plated on a Luria Broth (LB) agar plate containing 50 µg/ml kanamycin sulfate (Km), 20 µg/ml X-gal and 100 µM Isopropyl *ß*-D-1-thiogalacto-pyranoside (IPTG) for selection.

The plasmid pMM0092 was generated from PCR amplicons from two homologues regions (~500 bps) from *R. capsulatus* genomic DNA (Strain B10S) up- and downstream of *fdhABCD* that were amplified with the primers P7, P8, P9 and P10. Plasmid pK18mobSacB was linearized via PCR with the primers P5 and P6. The primers introduced terminal BsaI cutting sites with the required 4 bp overhangs. All amplicons were purified and assembled by Golden Gate cloning. The reaction mix was used to transform chemo-competent DH5α cells. For selection, the outgrowth was plated on LB agar plates containing 50 µg/ml Km, 20 µg/ml X-gal and 100 µM IPTG.

The plasmid pMM0205 was generated by introducing the *modABC* operon into pMM0106. The *modABC* gene was PCR amplified from genomic DNA (Strain B10S) with primers P11 and P12. The amplicon was purified and then cloned into plasmid pMM0106 (pK18mobSacB with BsaI cutting sites) by Golden Gate cloning. The reaction mix was used to transform DH5α cells. The outgrowth was plated on LB agar plates containing 50 µg/ml Km for selection.

To generate pMM0132, the *nifHDK* operon was PCR amplified from genomic DNA (B10S) using the primers P13 and P14. After purification, the amplicon was cloned into pOGG024-$Km^R$ by Golden Gate cloning. Chemo competent DH5α cells were transformed with the reaction mix and the outgrowth was plated on LB agar plates supplemented with 50 µg/ml Km for selection.

The plasmid pMM0190 was generated from pMM0132 by introducing an N-terminal 6x His-Tag at *anfH* and a C-terminal StrepTag II at *nifD* by restriction-free cloning using primers P15, P16, P17 and P18.[1]

### 3. *Rhodobacter capsulatus* strain construction

All strains used in this study are listed in Table 1. Genetic modifications of *Rhodobacter capsulatus* (i.e., the deletion of *fdhABCD* , the insertion of *modABC* and the insertion of *nifD*) were done with the *sacB* method described in [2]. In brief, plasmids for modification of the respective loci (pMM0092 for the deletion of *fdhABCD*, pMM0152 for the insertion of *modABC* and pMM0205 for the insertion of *nifD*) were introduced into the respective *R. capsulatus* strain via conjugation as described in [3], selecting for the kanamycin resistance conferred by the suicide vector. Single recombinant clones derived from single colonies of the previous step were passaged three times in liquid peptone yeast (PY) medium [3], growing each passage for 24 h at 30 °C and moderate shaking in the dark. The final passage was spread on a PY agar plate containing 5% (m/V) sucrose. The plate was incubated for 72 h at 30 °C under a closed atmosphere that was anaerobised using Oxoid™ AnaeroGen™ 3.5L sachets (Thermo Fisher Scientific Inc., Waltham, USA) and illumination by six 60 W krypton lamps (Osram Licht AG, Munich, Germany). Single colonies of *R. capsulatus* growing on the sucrose-containing agar plate were screened for kanamycin and sucrose sensitivity on PY plates containing 50 µg/mL kanamycin or 5% (m/V) sucrose, respectively. Colonies that could tolerate sucrose but were not growing on kanamycin-containing agar plates were further investigated via colony PCR to check the targeted genomic locus (*fdhABCD* deletion: P19, P20/21; *modABC* insertion: P22, P23/24; *nifD* insertion: P25, P26/27). Lastly, the purified PCR products were analyzed by Sanger sequencing (Microsynth Seqlab GmbH, Göttingen, Germany) to identify successful knockout clones. Plasmids for the recombinant expression of nitrogenase were introduced into *R. capsulatus* via conjugation, described in [3].

### 4. Protein production and purification

The Fe nitrogenase was purified according to established protocols.[4] For purification of the Mo nitrogenase, the *R. capsulatus* expression strain (MM0480) was inoculated on peptone yeast (PY) agar plates [3] containing 50 µg/mL kanamycin sulfate to select for the expression plasmid (MM0207). The plates were incubated phototrophically for 48 h at 32 °C under an Ar atmosphere. Obtained cell mass was used to inoculate liquid cultures in $N_2$-flushed RCV-Mo medium containing 30 mM DL-malic acid, 0.8 mM $MgSO_4$, 0.7 mM $CaCl_2$, 0.05 mM sodium ethylenediaminetetraacetic acid ($Na_2EDTA$), 0.03 mM thiamine hydrochloric acid, 9.4 mM $K_2HPO_4$, 11.6 mM $KH_2PO_4$, 5 mM serine, 45 µM $B(OH)_3$, 9.5 µM $MnSO_4$, 0.85 µM $ZnSO_4$, 0.15 µM $Cu(NO_3)_2$, 10 µM $Na_2MoO_4$ and 25 µg/mL kanamycin sulfate at a pH set to 6.8. The cultures were cultivated phototrophically at 32 °C for 24h. Subsequently, the liquid cultures were used to inoculate 800 mL of $N_2$-flushed RCV-Mo medium with an optical density of 0.1 at 660 nm ($OD_{660}$) for protein production. Protein purification was initiated when the cultures reached an $OD_{660}$ of ~ 3.0. Isolation of the Mo nitrogenase catalytic and reductase components was performed analogously to the purification procedure that we established previously for the Fe nitrogenase.[4]

### 5. SDS-PAGE analysis

For sodium dodecyl sulfate polyacrylamide gel electrophoresis (SDS-PAGE), protein samples were denatured at 98 °C in Pierce™ Lane Marker Reducing Sample Buffer (Thermo Fisher Scientific) for 10 min. The sample tubes were centrifuged (17,000×g, 1 min) and the supernatant was loaded on a 4–20% Mini-PROTEAN TGX Stain-Free Gel (Bio-Rad Laboratories, Inc.) including PageRuler™ Plus Prestained Protein Ladder (Thermo Fisher Scientific) as a molecular weight reference. The electrophoresis was run at a constant voltage of 180 V for 30 min. Bands were visualized by staining with GelCode™ Blue Safe Protein Stain (Thermo Fisher Scientific).

### 6. *In vitro* nitrogenase activity assays

Nitrogenase activity was assessed *in vitro* by measuring the specific activities for the formation of hydrogen ($H_2$), ammonia ($NH_3$), carbon monoxide (CO), methane ($CH_4$) and formate ($HCOO^-$) under varying atmospheres of dinitrogen ($N_2$), argon (Ar) and carbon dioxide ($CO_2$). Methods for the quantification of all products are described below. Assays were set up under an Ar atmosphere by dissolving the nitrogenase reductase component (molar excess over catalytic component as indicated) in an anaerobic solution of 50 mM TRIS (pH = 7.8), 10 mM sodium dithionite, 3.5 mM adenosine triphosphate (ATP), 7.87 mM $MgCl_2$, 44.59 mM creatine phosphate and 0.20 mg/mL creatine phosphokinase (catalog #C3755; Sigma-Aldrich St. Louis, USA). The reaction vials were sealed by crimping them with butyl rubber stoppers and the headspace was exchanged to 1.2 atm $N_2$, Ar or $CO_2$. For mixed atmospheres, indicated amounts of $CO_2$ were added to $N_2$ or Ar as indicated. Following a 10 min incubation at 30 °C, the reactions were initialized by adding 0.1 mg of nitrogenase catalytic component to a final volume of 700 µL. Reactions were allowed to proceed at 30 °C and moderate shaking at 250 rpm for 9 min before quenching with 300 µL 400 mM sodium ethylenediaminetetraacetic acid solution (pH = 8.0).

### 7. *In vivo* nitrogenase activity assays

*R. capsulatus* strains were inoculated from a glycerol stock on PY-agar plates [3] containing 20 µg/µl streptomycin sulfate (Sm). If the strain carried a nitrogenase expression plasmid, the agar contained 50 µg/mL kanamycin sulfate (Km) instead of Sm. Plates were incubated phototrophically for 48 h at 30 °C under an Ar atmosphere. Obtained cell mass was used to inoculate liquid cultures in $N_2$-flushed RCV minimal medium containing 30 mM DL-malic acid, 0.8 mM $MgSO_4$, 0.7 mM $CaCl_2$, 0.05 mM sodium ethylenediaminetetraacetic acid ($Na_2EDTA$), 0.03 mM thiamine hydrochloric acid, 9.4 mM $K_2HPO_4$, 11.6 mM $KH_2PO_4$, 120 µM $FeSO_4$, 45 µM $B(OH)_3$, 9.5 µM $MnSO_4$, 0.85 µM $ZnSO_4$, 0.15 µM $Cu(NO_3)_2$ at a pH set to 6.8. For *R. capsulatus* strains depending on the Mo nitrogenase for diazotrophic growth, 10 µM $N_2MoO_4$ were added. Subsequently, the headspace of the culture flasks was exchanged to 1.2 atm $N_2$ to allow for diazotrophic growth. For the complementation experiment with *R. capsulatus* strains MM0262 and MM0263 (Figure 5c), the RCV minimal medium was flushed with Ar instead of $N_2$ and was supplemented with 25 µg/mL kanamycin sulfate and 10 mM glutamine. Moreover, the culture headspace contained Ar instead of $N_2$. In all cases, the bacteria were cultivated phototrophically for 24 h at 30 °C. Subsequently, 10 mL cultures were inoculated at an $OD_{660}$ of 0.1 in the identical medium as the previous culture. The headspace was exchanged to 1.2 atm Ar, $N_2$ or 8 % $CO_2$ in Ar. For mixed atmospheres, $CO_2$ was added as indicated. The liquid cultures were cultivated phototrophically at 30 °C and samples for $HCOO^-$ quantification were taken at the indicated time points by piercing the septum of the cultivation vials with a $N_2$-flushed syringe.

### 8. Quantification of CO, $CH_4$ and $H_2$

Amounts of *in vivo* or *in vitro* formed CO, $CH_4$ and $H_2$ were determined via headspace analysis using a Clarus®690 GC system (GC–FID/TCD; PerkinElmer Inc., Waltham, USA) with a custom-made column circuit (ARNL6743) that was operated with the TotalChrom v. 6.3.4 software (PerkinElmer Inc., Waltham, USA). The headspace samples were injected by a TurboMatrixX110 (PerkinElmer Inc., Waltham, USA) autosampler, heating the samples to 45 °C for 15 min prior to injection. The samples were then separated on a HayeSep column (7’ HayeSep N 1/8’’ Sf; PerkinElmer Inc., Waltham, USA), followed by a molecular sieve (9’ Molecular Sieve 13x 1/8’’ Sf; PerkinElmer Inc., Waltham, USA) kept at 60 °C. Subsequently, the gases were detected with a flame ionization detector (FID, at 250 °C) and a thermal conductivity detector (TCD, at 200 °C). The quantification of all substrates was based on a linear standard curve derived from measuring varying amounts of CO, $CH_4$ and $H_2$ under identical conditions. The results were plotted using the GraphPad Prism 9 software (Dotmatics, Boston, USA).

### 9. Quantification of $NH_3$

Quantifying *in vitro* generated ammonia ($NH_3$) was done with a modified version of a fluorescence $NH_3$ quantification method described in [5]. 100 µL sample were combined with 1 mL of a solution containing 2 mM *o*-phthalaldehyde, 10 % (V/V) ethanol, 0.05 % (V/V) β-mercaptoethanol and 0.18 M potassium phosphate buffer (pH = 7.3) and incubated at 25 °C for 2 h in the dark. 50 µL of each sample were transferred into individual wells of a black Nunc™ F96 MicroWell™ plate (Thermo Fisher Scientific Inc., Waltham, USA) and fluorescence at 485 nm was monitored with an Infinite® 200 PRO plate reader (Tecan Group Ltd, Männedorf, Switzerland) in fluorescence top reading mode using an excitation wavelength of 405 nm. The quantification of ammonia was based on a linear standard curve that was derived from measuring varying amounts of $NH_4Cl$ under identical conditions. Samples incubated under an argon atmosphere instead of dinitrogen were used to correct for background signal. The results were plotted using the GraphPad Prism 9 software (Dotmatics, Boston, USA).

### 10. Quantification of formate via GC-MS

Formate quantification via gas chromatography-mass spectrometry (GC-MS) was performed according to [6]. In brief, 0.1 mL of sample was mixed with 0.5 mL 100 mM pentafluorobenzylbromide solution and 0.1 mL 500 mM potassium phosphate buffer (pH 6.8). The combined sample was mixed vigorously and then incubated for 1 h at 60 °C while shaking at 500 rpm. Samples were allowed to cool down to room temperature and 1 mL of a 100 µM 1,3,5-tribromobenzene solution in n-hexane was added. The samples were again mixed vigorously before separating the two phases by centrifugation (15 min, 1400 × g). 300 µL of the organic phase were transferred into a 1.5 ml short thread vial (VWR, Radnor, USA) and analyzed on a Clarus® 690 gas chromatograph (split = 1:20), equipped with an Elite-1 column that was coated with dimethylpolysiloxane (60 m, 0.32 mm i.d., 5.0 µm film thickness; PerkinElmer Inc., Waltham, USA). The gas chromatograph was connected to a Clarus® SQ8 T mass spectrometer (PerkinElmer Inc., Waltham, USA) and the system was operated with the TurboMass GC-MS Software ver. 6.1.2 (PerkinElmer Inc., Waltham, USA). The column's initial temperature (50 °C) was held for 3 min and then increased to 280 °C at a rate of 30 °C/min. The injection port and ion source were kept at 220 °C. Helium was used as the carrier gas at a flow rate of 1.5 mL/min. Mass spectra were obtained by positive-ion electron ionization (EI) mode scanning every 0.1 s from 40 – 600 *m/z*. Selected ion recording (SIR) was measured every 0.1 s for the molecular peak ion of the derivative of $HCOO^-$ at 226 *m/z* and the base peak ion of 1,3,5-tribromobenezene at 314 *m/z.* The ionization energy of the EI condition was 70 eV. A linear standard curve ($R^2 \geq 0.998$) for $HCOO^-$ was generated by plotting the ratio of the peak at 226 *m/z* and 314 *m/z* against the concentration of $HCOO^-$. The detection limit of $HCOO^-$ was 0.03 mM. The results were plotted using the GraphPad Prism 9 software (Dotmatics, Boston, USA).

### 11. Purification of formate dehydrogenase

Formate dehydrogenase (FDH) from *Pseudomonas* sp. 101 used for enzymatic $HCOO^-$ quantification was purified according to published protocols [7]. In brief, *E. coli* BL21 DE3 cells expressing N-terminally His-tagged FDH from plasmid pTE1390 were inoculated on LB agar plates containing 20 µg/mL of Sm. The plates were incubated at 37 °C for 24 h and then used to inoculate liquid cultures in terrific broth (TB) medium containing 20 µg/mL of Sm. Following 16 h of incubation at 30 °C and moderate shaking (80 rpm), the cells were harvested by centrifugation (15 min at 6000 × g, 10 °C) and then resuspended in 30 mL of binding buffer (20 mM Tris, 500 mM NaCl, 5 mM Imidazole, pH 7.9) per gram of pellet. Next, the cell suspension was supplemented with 0.2 mg/mL bovine pancreatic deoxyribonuclease I and incubated for 20 min on ice before lysing the cells by sonication. The lysate was centrifuged (20 min at 4347 × g, 4 °C) and the supernatant was loaded on 3 mL of Protino Ni-NTA agarose resin in a gravity flow column. Subsequently, the column was washed with 60 mL of washing buffer (20 mM, Tris, 500 mM NaCl, 20 mM imidazole, pH 7.9) before eluting the protein with elution buffer (20 mM, Tris, 500 mM NaCl, 250 mM imidazole, pH 7.9). The elution fraction was buffer exchanged to 100 mM $Na_2HPO_4$ buffer (pH 7.0) using an Amicon® Ultra-15 Centrifugal Filter Unit (molecular weight cut off = 30 kDa; Merck Millipore, Billerica, USA). The obtained protein solution was flash-frozen in liquid $N_2$ and stored at −70 °C until further use. Protein yields were determined using the Quick Start™ Bradford 1x Dye Reagent (Bio-Rad Laboratories, Inc., Hercules, USA) according to the instructions by the manufacturer and the purity was assessed via sodium dodecyl sulfate polyacrylamide gel electrophoresis (SDS-PAGE).

### 12. Quantification of formate via formate dehydrogenase assay

The FDH assay reaction mix contained 10 mM nicotinamide adenine dinucleotide ($NAD^+$) and 0.1 mg/mL purified FDH dissolved in 100 mM sodium phosphate buffer (pH 7.0). 97µl of the reaction mix was added to each well of a 96-well plate and absorbance at 340 nm ($A_{340}$) was monitored using a CLARIOstar Plus plate reader (shaking plate 30s at 400 rpm before each measurement, assay temperature set to 30 °C; BMG labtech, Ortenberg, Germany). Once $A_{340}$ remained stable, 3 µl of a sample was added to each well using a VIAFLO 96 pipetting station (Integra, Biebertal, Germany) and monitoring of $A_{340}$ was continued until it remained constant. The $A_{340}$ value was used to quantify $HCOO^-$ based on a standard dilution series of sodium formate ranging from 2.5 mM to 30 mM that was quantified identically. The results were plotted using the GraphPad Prism 9 software (Dotmatics, Boston, USA).

### 13. Diazotrophic growth curves

*R. capsulatus* strains were inoculated from a glycerol stock on PY-agar plates [3] containing 20 µg/µl Sm. Plates were incubated phototrophically for 48 h at 30 °C under an Ar atmosphere. Obtained cell mass was used to inoculate liquid cultures in 10 mL $N_2$-flushed RCV minimal medium containing 30 mM DL-malic acid, 0.8 mM $MgSO_4$, 0.7 mM $CaCl_2$, 0.05 mM sodium ethylenediaminetetraacetic acid ($Na_2EDTA$), 0.03 mM thiamine hydrochloric acid, 4.7 mM $K_2HPO_4$, 5.8 mM $KH_2PO_4$, 120 µM $FeSO_4$, 45 µM $B(OH)_3$, 9.5 µM $MnSO_4$, 0.85 µM $ZnSO_4$, 0.15 µM $Cu(NO_3)_2$ at a pH set to 6.8. For *R. capsulatus* strains depending on the Mo nitrogenase for diazotrophic growth, 10 µM $N_2MoO_4$ were added. The headspace of the culture flasks was exchanged to 1.2 atm $N_2$ and the bacteria were cultivated phototrophically for 48 h at 30°C. Subsequent inoculation steps were performed strictly anaerobically by working under an Ar atmosphere. The grown cultures were used to inoculate 50 mL of identical $N_2$-flushed RCV minimal medium at an $OD_{660}$ of 0.5, which were cultivated phototrophically under 1.2 atm $N_2$ atmosphere for 48 at 30 °C. The bacteria were then used to inoculate 6 mL of identical $N_2$-flushed RCV minimal medium in 22 mL glass vials at an $OD_{660}$ of 0.1. The headspace of the culture flasks was exchanged to 1.2 atm $N_2$ and cultivated phototrophically at 30 °C. Samples for $OD_{660}$ measurements were taken at the indicated time points by piercing the septum of the cultivation vials with an $N_2$-flushed syringe. The results were plotted using the GraphPad Prism 9 software (Dotmatics, Boston, USA). The doubling time $T_d$ was determined for the exponential growth phase by plotting ln($OD_{660}$/$OD_{660}$(t=0)) versus the time of growth. The data points were then interpolated by a linear regression model to determine the growth rate *r*. $T_d$ was calculated with equation (4):

$$T_d = \frac{\ln(2)}{r} \tag{4}$$

### 14. $^{13}CO_2$ labelling experiment

Cultures of *R. capsulatus* strains were prepared as described for the *in vivo* activity assays. Following inoculation at $OD_{660}$ of 0.1 and exchange of the culture headspace to $N_2$, 20 % (V/V) $^{13}CO_2$ (99.0% atom-% $^{13}C$) or $^{12}CO_2$ ($^{13}C$ at natural abundance) were added to the culture vial headspace. The cultures were cultivated phototrophically at 30 °C for six days. 5 mL of the cultures were centrifuged (4347 × g, 15 min) and 4 ml of the supernatant were lyophilized with an Alpha 1-4 LSCplus freeze dryer (Martin Christ Gefriertrocknungsanlagen GmbH, Osterode am Harz, Germany). The remaining solids were dissolved by vortexing in 1 ml $D_2O$ containing 8.33 mg/ml 3-(Trimethylsilyl)propionic-2,2,3,3-acid sodium salt D4 (TSPD4) as an internal standard. The solution was centrifuged (16000 × g, 10 min) and 0.6 mL were transferred into glass tubes. Nuclear magnetic resonance (NMR) spectra were measured at 298 K with an Avance II 300 MHz ($^1H$ 300 MHz) NMR spectrometer (Bruker, Billerica, USA). The spectra were calibrated to the residual signal of HOD ($\delta$ = 4.79 ppm). The signal intensity of TSP-D4 ($I_{TPS\text{-}D4}$) was used for the quantification of $HCOO^-$ according to equation (5):

$$N_{HCOO\text{-}} = \frac{I_{HCOO\text{-}} \cdot 9}{I_{TPSD-D4}} \cdot N_{TPS-D4} \tag{5}$$

The quantification was validated by generating a standard curve for $HCOO^-$ measuring the amount of $HCOO^-$ in samples with known concentrations. The detection limit for $HCOO^-$ was 0.6 mM. The $^{13}C/^{12}C$-ratio of $HCOO^-$ in the culture supernatant was derived by dividing the NMR peak integral of $H^{13}COO^-$ ($^1H$ NMR (300.0 MHz, $D_2O$): $\delta$ 8.44 (d, $^1J(H,^{13}C)$ = 195 Hz, 1H) ppm) by the peak integral of $H^{12}COO^-$ ($^1H$ NMR (300.0 MHz, $D_2O$): $\delta$ 8.44 (s, 1H) ppm).

### 15. Supporting tables

#### Table 1. Strains used in this study.

| Strain | Relevant characteristics | Reference |
|---|---|---|
| *R. capsulatus* BS85 | Polar *nifD*::$Sp^R$ mutant (Δ*nifDK*) of B10S; $Sm^R$, $Sp^R$ | Hoffmann *et al.*[8] |
| *E. coli* ST18 | RP4-2 *Tc*::*Mu Km*::*Tn7* Δ*hemA* mutant | Thoma *et al.*[9] |
| *E. coli* DH5α | General cloning strain | NEB® Biolabs |
| *E. coli* BL21 DE3 | T7 Expression Strain | NEB® Biolabs |
| *R. capsulatus* MM0425 | Δ*anfHDGK*::*gmR* Δ*modABC* Δ*draTG* Δ*gtaI* Δ*nifHDK;* $Sm^R$, $Sp^R$, $Gm^R$ | Schmidt *et al.*[4] |
| *R. capsulatus* MM0437 | MM0425 carrying pMM0128; $Sm^R$, $Sp^R$, $Km^R$ | Schmidt *et al.*[4] |
| *R. capsulatus* MM0468 | modABC insertion mutant of MM0425*;* $Sm^R$, $Sp^R$, $Gm^R$ | This work |
| *R. capsulatus* MM0480 | MM0468 carrying pMM0207; $Sm^R$, $Sp^R$, $Gm^R$, $Km^R$ | This work |
| *R. capsulatus* MM0335 | WT*, Sp*::*nifD* of BS85; $Sm^R$, used in Figure 4a, Figure 4b, Figure 5a and Figure 5e | This work |
| *R. capsulatus* MM0057 | Δ*modABC* mutant of BS85; $Sm^R$, $Sp^R$, used in Figure 4c, Figure 4d, Figure 5a, Figure 5b, Figure 5c and Figure 5e | Schmidt *et al.*[4] |
| *R. capsulatus* MM0372 | Δ*fdhABCD* mutant of MM0335; $Sm^R$, used in Figure S4 and Figure S5 | This work |
| *R. capsulatus* MM0302 | Δ*fdhABCD* mutant of MM0057; $Sm^R$, $Sp^R$, used in Figure S4 and Figure S5 | This work |
| *R. capsulatus* MM0164 | Polar *anfHDGK::Gm^R* mutant of MM00302; $Sm^R$, $Sp^R$, $Gm^R$ | This work |
| *R. capsulatus* MM0262 | MM0164 carrying pOGG024-$Km^R$; $Sm^R$, $Sp^R$, $Gm^R$, $Km^R$,used in Figure 5d | |
| *R. capsulatus* MM0263 | MM0164 carrying pMM0128; $Sm^R$, $Sp^R$, $Gm^R$, $Km^R$, used in Figure 5d | This work |

**Table 2: Plasmids used in this study.**

| | | |
|---|---|---|
| pK18*mobSacB* | Suicide vector, mobilizable (*oriT*), *sacB,* $Km^R$ | Schafer *et al.*[2] |
| pMM0106 | pK18mobSacB variant with a BsaI golden gate cloning site; $Km^R$ | This work |
| pOGG024 | Broad host range plasmid, mobilizable (*oriT*), *lacZα* cassette for golden gate cloning (BsaI), $Gm^R$ | Geddes *et al.*[10] |
| pOGG024-$Km^R$ | $Gm^R$ in pOGG024 exchanged to $Km^R$ | Schmidt *et al.*[4] |
| pMM0156 | To reintroduce *nifD*. Derived from pK18*mobSacB* | This work |
| pMM0205 | To reintroduce *modABC*. Derived from pMM0106 | This work |
| pMM0092 | To delete *fdhABCD*. Derived from pK18*mobSacB* | This work |
| pMM0128 | pOGG024-$Km^R$ *anfHDGK* expression plasmid (natural promotor) | Schmidt *et al.*[4] |
| pMM0190 | pOGG024-$Km^R$ *anfHDGK* expression plasmid (natural promotor) N-terminal 6x His-Tag *anfH*; C-terminal Strep-Tag II *anfD* | Schmidt *et al.*[4] |
| pMM0132 | pOGG024-$Km^R$ *nifHDK* expression plasmid (natural promotor) | This work |
| pMM0207 | pOGG024-$Km^R$ *nifHDK* expression plasmid (natural promotor) N-terminal 6x His-Tag *nifH*; C-terminal Strep-Tag II *nifD* | This work |
| pTE1390 | Formate dehydrogenase from Pseudomonas sp. 101 N-terminal 6x His-Tag, $Sm^R$ | Calzadiaz-Ramirez *et al.*[7] |

**Table 3: Primers used in this study.**

| Primer | Sequence | Used for |
|---|---|---|
| P1 (oMM0079) | CGTAATAGCGAAGAGGCCCG | Construction of pMM0156 (to insert *nifD*) |
| P2 (oMM0080) | GTAAAACGACGGCCAGTGC | |
| P3 (oMM0506) | GCACTGGCCGTCGTTTTACTCAGCGGGTCAGATCGAAGC | |
| P4 (oMM0507) | GGGCCTCTTCGCTATTACGGGCGGGGTTTTTCATTTCCTC | |
| P26 (oMM0130) | AGCAGTGGATCAGGTTCAGC | Colony PCR of the *nifD* genomic region |
| P25-(oMM0350) | ACCCATCTCGACAACAAGCC | |
| P27 (oMM0420) | TGCGTTCGGTCAAGGTTCTG | |
| P5 (oMM0227) | GATTTAGGTCTCTGTAAAACGACGGCCAGTGC | Construction of pMM0092 (to delete *fdhABCD*) |
| P6 (oMM0228) | GATTTAGGTCTCTCGTAATAGCGAAGAGGCCCG | |
| P7 (oMM0229) | GATTTAGGTCTCTTTACGGATCTCGACATGCTGGTGG | |
| P8 (oMM0230) | GATTTAGGTCTCTAGACGATCCGCTGGTCATCGCC | |
| P9 (oMM0231) | GATTTAGGTCTCTGTCTTCGGGAATGAAACCAAAGG | |
| P10 (oMM0232) | GATTTAGGTCTCTTACGGAGATCGAGCATTACCCCGG | |
| P19 (oMM0281) | GGTCTCGGTGTCATACTGCG | Colony PCR of the *fdhABCD* genomic region |
| P20 (oMM0282) | GGTTTTTGACGGGGTTACGC | |
| P21 (oMM0283) | TCGTCTATTTCGCCTGGGTG | |
| P11 (oMM0699) | GATTTAGGTCTCTATGGAGTCTGACGATGCGCACTTC | pMM0205 (to introduce *modABC*) |
| P12 (oMM0700) | GATTTAGGTCTCTGGAGGCAGAACCGAATCCGAAAGC | |
| P13 (oMM0406) | GATTTAGGTCTCAGGAGTCAGCGGGTCAGATCGAAGC | pMM0132 (*nifHDK* expression) |
| P14 (oMM0407) | GATTTAGGTCTCTAGCGGGCGGGGTTTTTCATTTCCTC | |
| P15 (oMM0681) | GCGATCTGACGGAGTTTGCCATGGTGATGATGGTGGTG | Construction of pMM0190 (*nifHDK* expression) |
| P16 (oMM0682) | CCCAAGGGAGCCACACATGCACCACCATCATCACCAT | |
| P17 (oMM0683) | GCCCCCCTTGGGCTCATTTTTCGAACTGCGGGTGG | |
| P18 (oMM0684) | CCATCGCCGCCGAGTGGAGCCACCCGCAGTTCG | |

### 16. Supporting figures

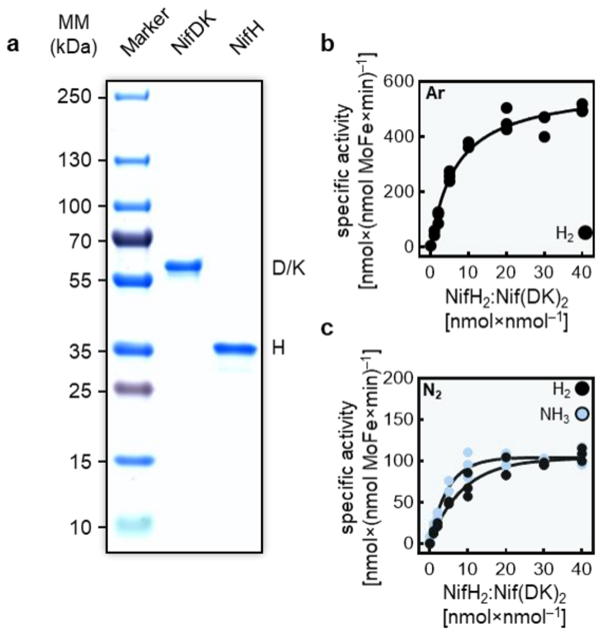

**Figure S1**. **Purification and biochemical characterization of the Mo nitrogenase.** (**a**) SDS-PAGE analysis of the purified Mo nitrogenase catalytic component (NifDK) and reductase component (NifH). (**b, c**) $NH_3$ and $H_2$ formation rates of the Mo nitrogenase under 1.2 atm atmospheres of Ar or $N_2$. Each assay was conducted with 100 µg of catalytic component with a varying molar excess of reductase component. Dots represent individual values of three replicates (n = 3).

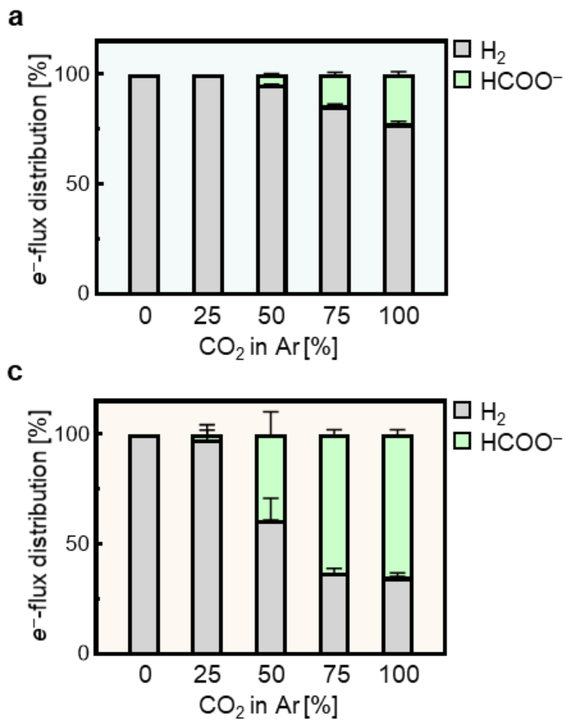

**Figure S2. Electron flux analysis of $CO_2$ reduction assays of the Mo and Fe nitrogenase in argon.** Electron flux distribution for *in vitro* activity assays of Mo (**a**, **b**) and Fe nitrogenase (**c**, **d**) under increasing concentrations of $CO_2$ in Ar. Bars and error bars represent mean ± s.d. from replicates (n = 3).

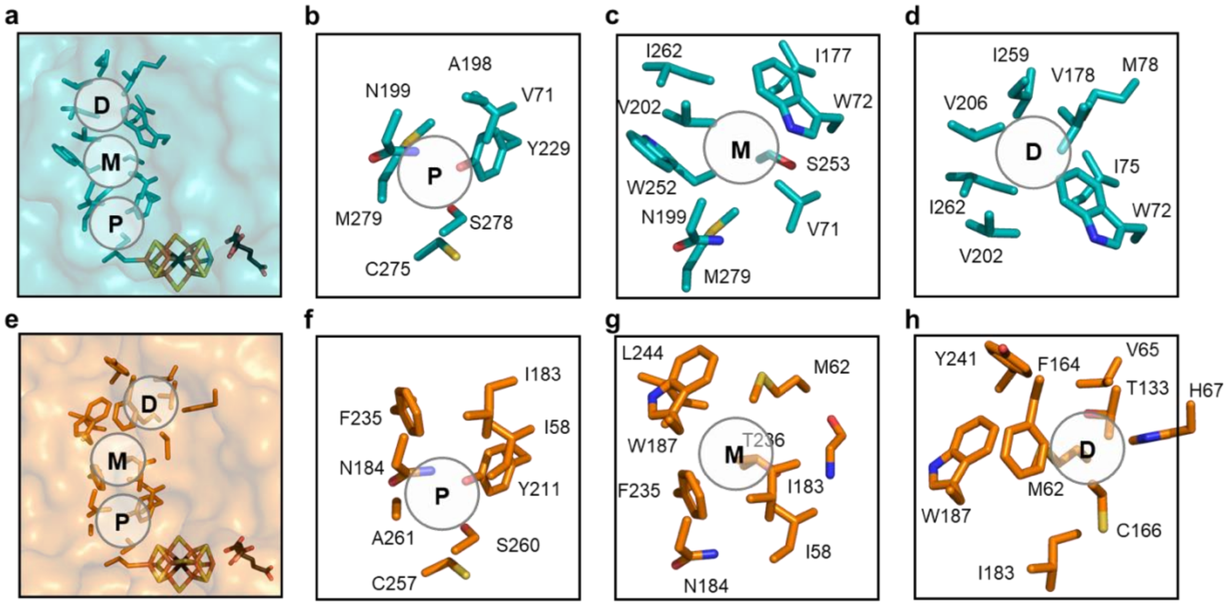

**Figure S3: The Igarashi-Seefeldt nitrogenase substrate channel.** (**a**) Surface representation of AnfD from the *Azotobacter vinelandii* Mo nitrogenase (PDB: 6UG0) with residues of the Igarashi-Seefeldt (IS) channel and the FeMoco displayed as sticks. (**b – d**) Focus on the proximal pocket (P), the medial (M) and the distal pocket (D) of the Mo nitrogenase as described by Gee *et al.*[11] (**e**) Surface representation of AnfD from the *R. capsulatus* Fe nitrogenase (PDB: 8OIE) with residues of the homologous Fe nitrogenase channel and the FeFeco displayed as sticks. (**f** – **h**) Focus on the proximal (P), medial (M) and distal (D) pocket of the homologous channel of the Fe nitrogenase, as identified by a structural comparison of NifD and AnfD. Four of the 15 channel residues of NifD are conserved for the Fe nitrogenase (C275, S278, Y229, N199), all located in the P-pocket. $M279^{NifD}$ of the Mo nitrogenase P-pocket is replaced by $A261^{AnfD}$ in the Fe isoform, causing $F235^{AnfD}$ (the homolog of $W252^{NifD}$) to participate in the P-pocket as well. In the M-pocket, $W252^{NifD}$ has recently been described to exhibit different conformations under turnover that could alter the IS channel access to the FeMoco.[12] The change of $W252^{NifD}$ to $F235^{AnfD}$ emphasizes the differences in substrate channeling between the two isoenzymes. Moreover, the residue $F164^{AnfD}$ is located at the position of the Mo nitrogenase D pocket, which causes a shift of the proposed D pocket of AnfD. In conclusion, the Mo nitrogenase IS channel and its hydrophobic character seem to be conserved in the Fe nitrogenase. Nevertheless, the AnfD channel architecture differs significantly from its NifD counterpart, potentially providing an explanation for the observed differences in $N_2$ versus $CO_2$ selectivity of Mo and Fe nitrogenases (Figure 3).

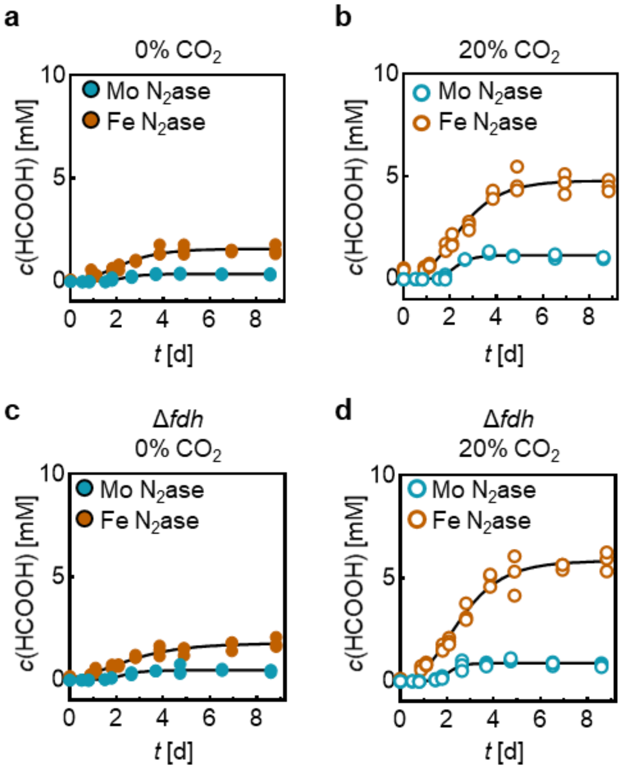

**Figure S4: *In vivo* $CO_2$ reduction by nitrogenases.** (**a**) Accumulation of $HCOO^-$ over time in the culture medium of diazotrophically growing *R. capsulatus* strains depending on either the Mo or the Fe nitrogenase. (**b**) Identical experiment as (a) but with 20% $CO_2$ added to the $N_2$ in the headspace. (**c**, **d**) Identical experiments to (a) and (b) but strains carrying an additional deletion of the formate dehydrogenase (Δ*fdh*). Dots represent values from each replicate (n = 3).

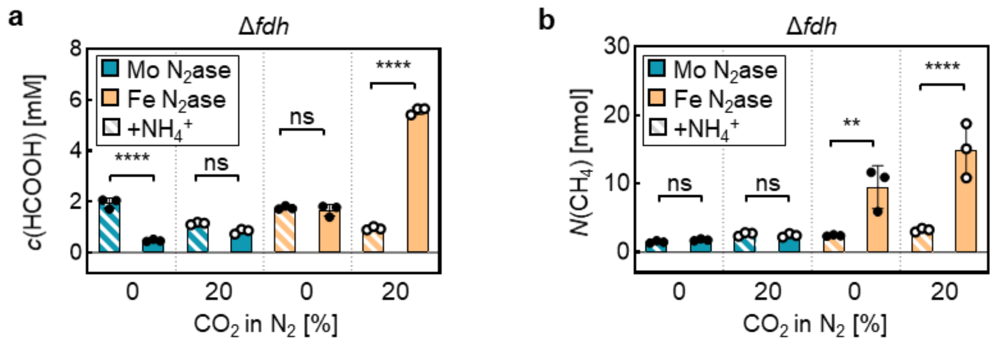

**Figure S5: *In vivo* $CO_2$ reduction by nitrogenase in formate dehydrogenase deletion (Δ*fdh*) *R. capsulatus* strains.** (**a**) $HCOO^-$ concentration in *R. capsulatus* culture supernatant expressing the Mo nitrogenase (MM0372) or the Fe nitrogenase (MM0302) under diazotrophic growth conditions in the absence or presence of 20% $CO_2$ after 6 d of growth. (**b**) Amount of $CH_4$ measured in the culture headspace of *R. capsulatus* cells after 9 d of growth determined by GC-FID. Dots represent values from biological replicates (n = 3).

### 17. Data availability statement

All unique materials used in this study are available from the corresponding author upon request. All raw data for *in vivo* work, kinetic experiments and protein characterizations will be deposited on Edmond, the Open Research Data Repository of the Max Planck Society and available upon publication
